# HIV-1 Protease Evolvability is Affected by Synonymous Nucleotide Recoding

**DOI:** 10.1101/315366

**Authors:** Maria Nevot, Ana Jordan-Paiz, Glòria Martrus, Cristina Andrés, Damir García-Cehic, Josep Gregori, Sandra Franco, Josep Quer, Miguel Angel Martinez

## Abstract

One unexplored aspect of HIV-1 genetic architecture is how codon choice influences population diversity and evolvability. Here we compared the development of HIV-1 resistance to protease inhibitors (PIs) between wild-type (WT) virus and a synthetic virus (MAX) carrying a codon-pair re-engineered protease sequence including 38 (13%) synonymous mutations. WT and MAX viruses showed indistinguishable replication in MT-4 cells or PBMCs. Both viruses were subjected to serial passages in MT-4 cells with selective pressure from the PIs atazanavir (ATV) and darunavir (DRV). After 32 successive passages, both the WT and MAX viruses developed phenotypic resistance to PIs (IC_50_ 14.6 ± 5.3 and 21.2 ± 9 nM for ATV, and 5. 9 ± 1.0 and 9.3 ± 1.9 for DRV, respectively). Ultra-deep sequence clonal analysis revealed that both viruses harbored previously described resistance mutations to ATV and DRV. However, the WT and MAX virus proteases showed different resistance variant repertoires, with the G16E and V77I substitutions observed only in WT, and the L33F, S37P, G48L, Q58E/K, and L89I substitutions detected only in MAX. Remarkably, G48L and L89I are rarely found *in vivo* in PI-treated patients. The MAX virus showed significantly higher nucleotide and amino acid diversity of the propagated viruses with and without PIs (*P* < 0.0001), suggesting higher selective pressure for change in this recoded virus. Our results indicate that HIV-1 protease position in sequence space delineates the evolution of its mutant spectra. Nevertheless, the investigated synonymously recoded variant showed mutational robustness and evolvability similar to the WT virus.

**IMPORTANCE:** Large-scale synonymous recoding of virus genomes is a new tool for exploring various aspects of virus biology. Synonymous virus genome recoding can be used to investigate how a virus’s position in sequence space defines its mutant spectrum, evolutionary trajectory, and pathogenesis. In this study, we evaluated how synonymous recoding of the human immunodeficiency virus type 1 (HIV-1) protease impacts the development of protease inhibitor (PI) resistance. HIV-1 protease is a main target of current antiretroviral therapies. Our present results demonstrate that the wild-type (WT) virus and the virus with the recoded protease exhibited different patterns of resistance mutations after PI treatment. Nevertheless, the developed PI resistance phenotype was indistinguishable between the recoded virus and the WT virus, suggesting that the synonymously recoded protease HIV-1 and the WT protease virus were equally robust and evolvable.

## INTRODUCTION

Alterations in a DNA or mRNA sequence that do not change the protein amino acid sequence are called synonymous mutations. Although they do not influence the resulting protein sequence, synonymous mutations can still substantially affect cellular processes (1, 2). Notably, synonymous virus genome recoding can impact viral replication capacity and fitness (3), reportedly leading to attenuation of multiple RNA and DNA viruses, including poliovirus (4-7), influenza virus (8, 9), HIV-1 (10-12), SIV (13), Chikungunya virus (14), human respiratory syncytial virus (15-17), porcine reproductive and respiratory syndrome virus (18), echovirus 7 (19, 20), tick-borne encephalitis virus (21), vesicular stomatitis virus, dengue virus (22), adeno-associated virus (23), and papillomavirus (24).

Synonymous virus genome recoding is being investigated as a new strategy for generating novel live-attenuated vaccine candidates. This method is promising because the amino acid coding is completely unaffected, thereby avoiding the potential generation of new and undesirable biological properties. Moreover, synonymous virus genome recoding involves the introduction of hundreds or thousands of nucleotide substitutions, which minimizes the risk of phenotypic reversion via point mutations or through recombination with homologous sequences in circulating strains. This is particularly important with regards to RNA viruses, since viral RNA polymerases lack error-correction mechanisms (25-27). The high genetic variability of RNA viruses is a critical limitation when designing novel antiviral strategies.

The usefulness of synonymous virus genome recoding goes beyond the generation of new live attenuated vaccines. This method has also been used to identify specific RNA structures required for virus replication (28), virus genome cis-inhibitory signal sequences important for complex viral functions (23), and novel antiviral mechanisms within the innate immune response (11, 29, 30), as well as to resolve the importance of codon usage in the temporal regulation of viral gene expression (31). In one interesting example, synonymous virus genome recoding was used to demonstrate that a synonymous position in sequence space can impact poliovirus evolvability and pathogenesis (32, 33).

Like other RNA viruses, human immunodeficiency virus type 1 (HIV-1) populations comprise a closely related mutant spectra or mutant clouds termed viral quasispecies (34, 35). Mutant cloud composition can impact virus evolvability, fitness, and virulence (25-27). One unexplored aspect of HIV-1 genetic architecture is how codon choice influences population diversity and evolvability. It is presently unclear whether HIV-1 sequences have evolved to optimize both protein coding and DNA/RNA sequences. The HIV-1 genome exhibits a particularly striking bias towards enrichment of A-rich codons, which may be a selectable trait (36) and affect innate immune recognition (11, 29). Similarly, synonymous codon usage can temporally regulate expressions of structural gene products of the simian immunodeficiency virus (SIV) (31) and regulate HIV-1 splicing and replication (12).

Here we aimed to explore whether synonymous sequence space influences the development of protease inhibitor (PI) resistance, and to thus determine whether HIV-1 evolvability is influenced by the natural position of a protease in sequence space. To this end, we compared the development of HIV-1 resistance to the protease inhibitors atazanavir (ATV) and darunavir (DRV) between wild-type HIV-1 (WT) and synthetic HIV-1 carrying a synonymously recoded protease sequence (MAX).

## RESULTS

WT HIV-1 was compared with the MAX variant that carried a protease gene including 38 synonymous mutations (13% of the protease sequence) (Fig. 1). These 38 synonymous substitutions were scattered throughout the protease coding region, excluding the first 40 amino-terminal nucleotides that overlap with the carboxy-terminal of the gag p6 reading frame. These 38 substitutions were chosen in order to improve protease gene codon pair bias (6) without modifying its codon bias or folding free energy (10). The WT and MAX viruses have identical consensus amino acid sequences,-and, as we have previously demonstrated (10), this MAX variant and the WT virus show indistinguishable replication in MT-4 cells (Fig. 2) or PBMCs (10).

**FIG 1.**
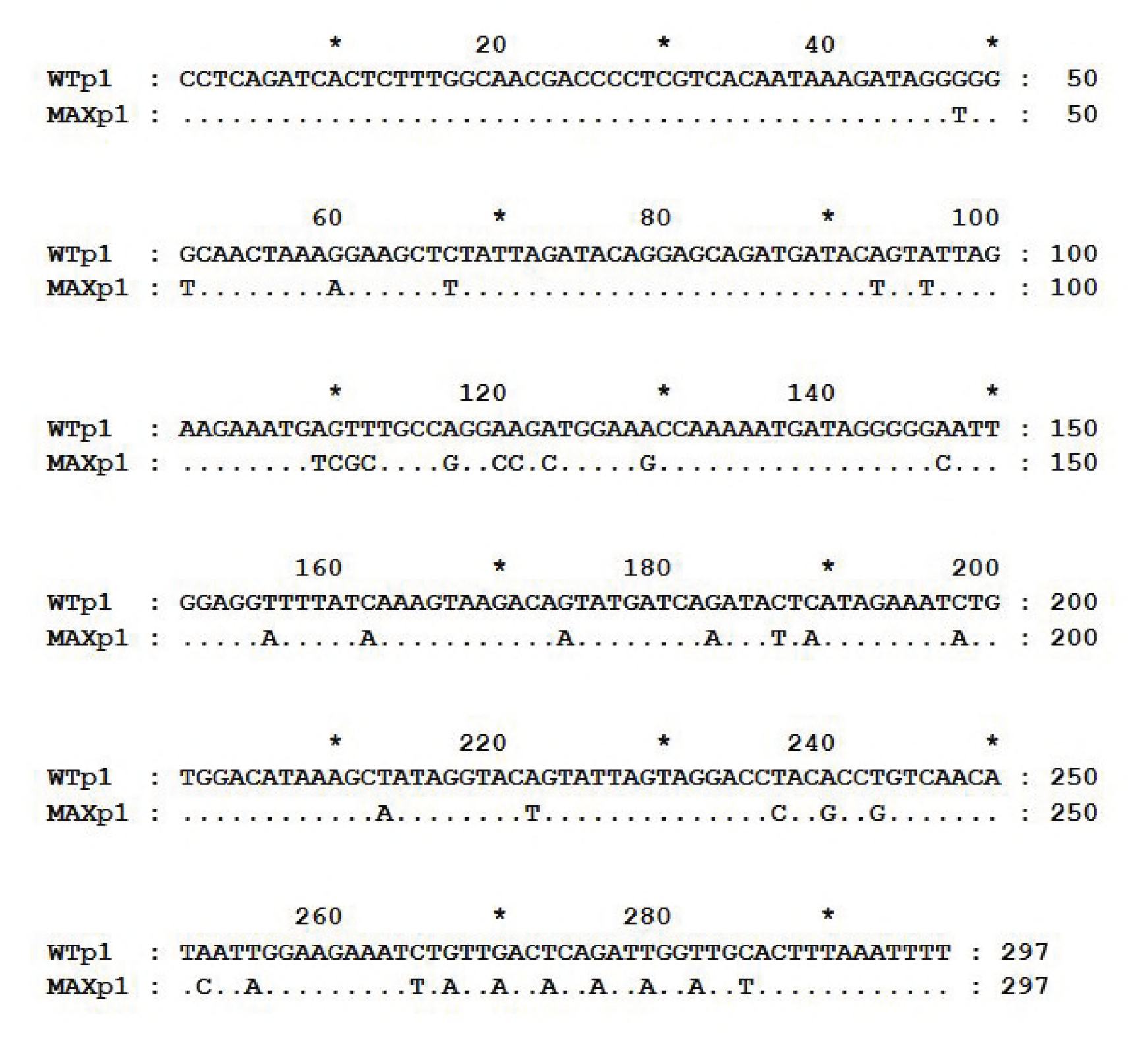
HIV-1 protease nucleotide sequences of the wild-type (WT) virus that corresponds to the HIV-1 HXB2 strain (http://www.hiv.lanl.gov), and of the synthetic MAX variant that was generated by PCR combining three overlapping synthetic DNA oligonucleotides as previously described (10). No substitution was introduced in the first 40 protease amino-terminal nucleotides that overlap with the carboxy-terminal of the gag p6 reading frame.

**FIG 2.**
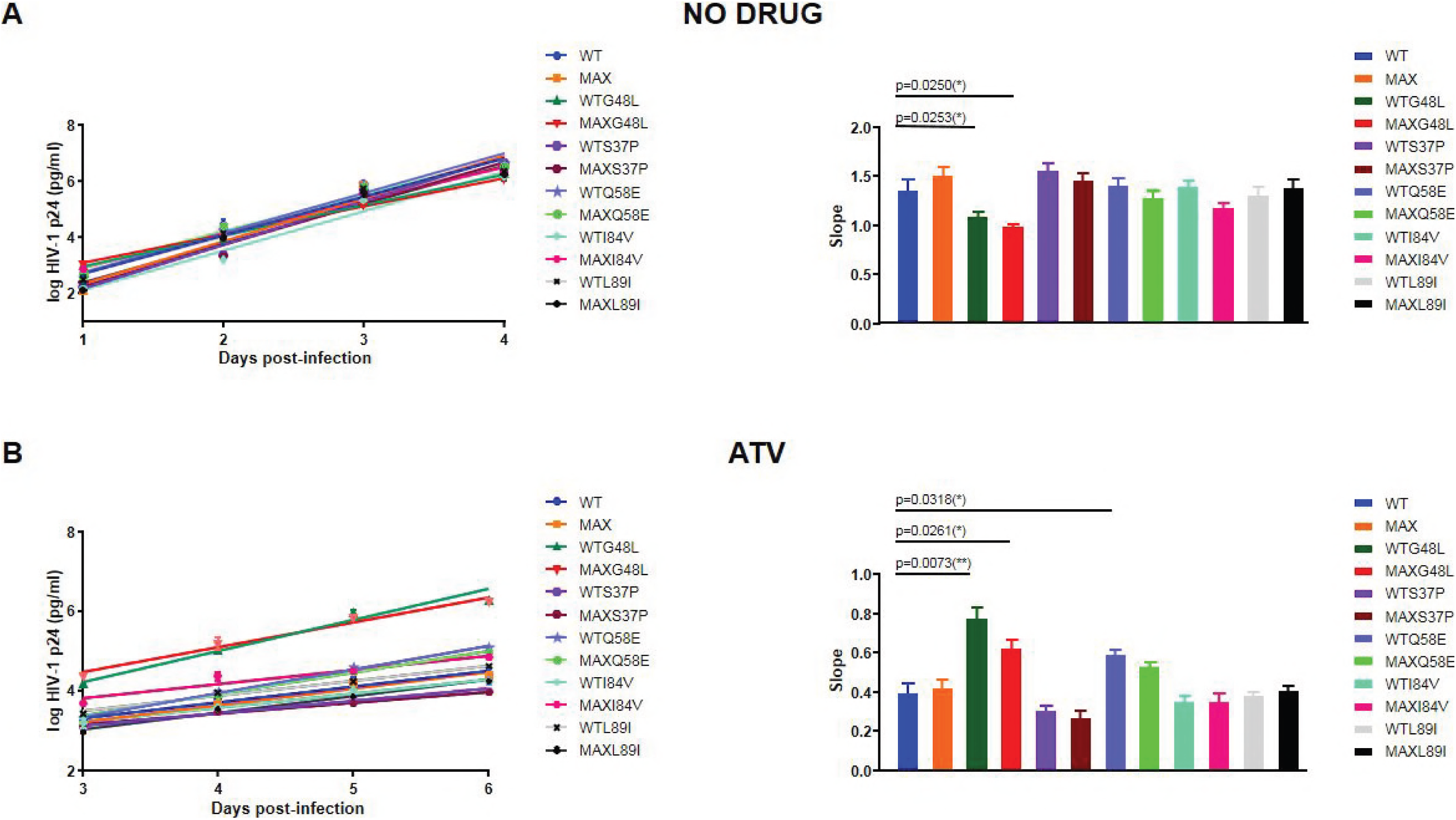
Replication kinetic assay of wild-type HIV-1 (WTp1) and the recoded MAX protease variant (MAXp1) in MT-4 cells. HIV-1 antigen p24 concentrations in culture supernatants were measured on days 0-4 in the absence of drug or days 0–6 in the presence of drug. For each virus, the slope of the plot provides an estimate of the viral replication capacity. Bars show the slope of the p24 antigen production from each virus after infection of MT-4 cells. Comparison between WTp1 (HXB2) and the mutant MAX recoded viruses are shown, as well as between the corresponding WT and MAX virus variants (i.e.. S37P, G48L, Q58E, I84V and I89L). The significance of the difference between slopes was calculated using an unpaired t-test with Whech’s correction in GraphPrism v. 7 software. **(A)** Kinetic assays performed in the absence of drug. All slopes values were statistically tested against the WT value. Only WTG48L and MAXG48L displayed a lower replication capacity than the WT. **(B)** Kinetic assays performed in the presence of 20nM atazanavir (ATV). All slopes values were statistically tested against the WT value. WTG48L, MAXG48L and WTQ58E displayed a higher replication capacity than the WT in the presence of ATV. Values represent the mean ± standard deviation (SD) from at least three independent experiments.

We subjected the WT and MAX viruses to selective pressure from two PIs: ATV and DRV. The viruses were propagated in duplicate in MT-4 cells over 32 serial passages (128 days of culture) without drugs or with increasing concentrations of ATV or DRV. The starting PI concentrations were near the half maximal inhibitory concentration (IC_50_) for the WT HXB2 virus (10). Before the passages, both viruses showed similar IC_50_ values for ATV and DRV (Table 1) (10). The WTp32 and MAXp32 viruses still showed comparable IC_50_ values after 32 serial passages in the presence or absence of PIs (Table 1). WTp32 and MAXp32, respectively, showed 5- fold and 13-fold increases in IC_50_ for ATV, and 6-fold and 10-fold increases in IC_50_ for DRV (Table 1). Although MAXp32 displayed a higher resistance to ATV and DRV than WTp32 (Table 1), these differences were not significant (P = 0.4816 and P = 0.3451, respectively). These assays demonstrated that the MAX variant virus did not show impaired capacity to develop phenotypic resistance to PIs.

**TABLE 1.** Susceptibility of HIV-1 carrying WT or MAX proteases to atazanavir (ATV) or darunavir (DRV)

| Protease | IC<sub>50</sub> (nM) |  |
| --- | --- | --- |
|  | ATV | DRV |
| WTp1 | 3.1 ± 1.2 (1)<sup>1</sup> | 0.9 ± 0.3 (1) |
| WTp32 | 14.6 ± 5.3 (5) | 5.7 ± 1.0 (6) |
| MAXp1 | 1.6 ± 0.1 (1) | 0.9 ± 0.2 (1) |
| MAXp32 | 21.2 ± 9.0 (13) | 9.3 ± 1.9 (10) |

<sup>1</sup>Fold change.

After the 32 cell passages in the presence of ATV or DRV, virus RNA was recovered. This RNA was RT-PCR amplified and ultra-deep sequenced, and we compared the frequencies of resistant mutations. For each of the two studied viruses and the two tested drugs, we sequenced between 1.9 × 10^7^ and 4.1 × 10^7^ individual protease nucleotides (Table 2). Sequence clonal analysis revealed no resistance-associated substitutions in viruses propagated without drugs (Table 3). On the other hand, both WT and MAX viruses propagated in the presence of PIs developed previously described resistance mutations to ATV and DRV (Table 3). Moreover, the resistance variant repertoire differed between the MAX and WT viruses (Table 3). Specifically, the G16E substitution was observed only in the WT protease virus propagated with ATV or DRV. Notably, the WT protease required only a transition to develop this substitution, whereas the MAX protease would need two substitutions, a transition, and a transversion. Additionally, the L33F, G48L, Q58E/K, and I89L substitutions were detected only in the recoded MAX protease. Other accompanying substitutions were also detected only in the MAX virus (e.g., E21K, H69Y, and T91S), mainly in the MAX virus (e.g., L10F), or only in the WT virus (e.g., L23I, P39Q, and V77I). Interestingly, some resistance mutations selected by the MAX protease virus (i.e., G48L and I89L), are extremely rare non-polymorphic substitutions *in vivo* (37). This finding indicates that the MAX protease may explore a different sequence space than that of the WT protease. Similar to G16E, the I89L mutation requires two substitutions in the WT background and only one substitution in the MAX background. However, there were no obvious reasons for the preferential emergence of K45I, G48L, Q58E and I84V in the MAX background. Substitutions L10F, G16E, L33F, Q58E/K, V77I, I84V, and I89L have been previously associated with PIs resistance (38). In contrast, E21K, L23I, P39Q, G48L, H69Y, and T91S have not been described as associated to PIs resistance.

**TABLE 2.** Substitutions associated with protease inhibitor resistance detected after MT-4 cell passages in the presence of atazanavir (ATV), darunavir (DRV), or no drug (ND)

| ATV | Experiment 1 |  | Experiment 2 |  | DRV | Experiment 1 |  | Experiment 2 |  | ND | Experiment 1 |  |
| --- | --- | --- | --- | --- | --- | --- | --- | --- | --- | --- | --- | --- |
|  | WTp1 | MAXp1 | WTp1 | MAXp1 |  | WTp1 | MAXp1 | WTp1 | MAXp1 |  | WTp1 | MAXp1 |
| L10F<sup>1</sup> | 1.0<sup>2</sup> | 28.0 | 0 | 0 | L10F | 0 | 19.2 | 0 | 0 | D30N | 1.4 | 2.3 |
| G16E | 1.7 | 0 | 1.8 | 0 | G16E | 16.5 | 0 | 55.5 | 0 | S37P | 0 | 11.7 |
| L23I | 0.8 | 0 | 0 | 0 | E21K | 0 | 1.4 | 0 | 0 | P39Q | 2.1 | 0 |
| V32A | 76.3 | 0 | 53.3 | 59.9 | A28S | 46.1 | 45.4 | 6.6 | 0 | P39T | 0.8 | 0 |
| V32I | 10.1 | 24.0 | 2.9 | 23.8 | L33F | 0 | 94.2 | 0 | 89.2 | R41K | 0.8 | 0 |
| V32T | 0.65 | 0 | 0 | 0 | P39Q | 0.6 | 0 | 0.6 | 0 | K43R | 0 | 0.5 |
| L33F | 0 | 0.8 | 0 | 0 | K45I | 0 | 0 | 0 | 9.8 | Q58K | 0 | 1.1 |
| P39Q | 0.9 | 0 | 0.9 | 0 | M46I | 6.0 | 3.9 | 94.0 | 14.6 | I72T | 1.8 | 0 |
| K45I | 0 | 10.2 | 6.0 | 0 | I50L | 49.7 | 52.3 | 39.1 | 100 | V82I | 1.3 | 0 |
| K45R | 0 | 0.8 | 0 | 0 | Q58K | 0 | 0 | 0 | 1.2 |  |  |  |
| M46I | 12.1 | 0 | 1.1 | 10.4 | H69Y | 0 | 0.6 | 0 | 0 |  |  |  |
| G48L | 0 | 0 | 0 | 22.4 | A71V | 0 | 25.2 | 0 | 0 |  |  |  |
| I50L | 0 | 28.6 | 27.3 | 0 | V82I | 85.2 | 0.6 | 34.8 | 19.7 |  |  |  |
| Q58E | 0 | 20.2 | 0 | 0 | I84V | 3.4 | 0 | 0 | 0 |  |  |  |
| Q58K | 0 | 0 | 0 | 0.8 |  |  |  |  |  |  |  |  |
| A71V | 88.2 | 23.2 | 62.8 | 71.1 |  |  |  |  |  |  |  |  |
| V77I | 0 | 0 | 11.4 | 0 |  |  |  |  |  |  |  |  |
| V82I | 0 | 0 | 9.8 | 0 |  |  |  |  |  |  |  |  |
| V82D | 0 | 0 | 0 | 1.2 |  |  |  |  |  |  |  |  |
| I84V | 0 | 43.0 | 0 | 0 |  |  |  |  |  |  |  |  |
| N88S | 14.0 | 28.2 | 19.7 | 3.4 |  |  |  |  |  |  |  |  |
| I89L | 0 | 0 | 0 | 59.4 |  |  |  |  |  |  |  |  |
| T91S | 0 | 3.6 | 0 | 0 |  |  |  |  |  |  |  |  |

<sup>1</sup>HIV-1 protease amino acid position.
<sup>2</sup>Percentage of the corresponding substitution.

**TABLE 3.** Susceptibility of HIV-1 carrying WT or MAX protease variants to atazanavir (ATV).

| Protease | ATV IC<sub>50</sub> (nM) |
| --- | --- |
| WT | 1.8 ± 1.2 (1)<sup>1</sup> |
| MAX | 1.6 ± 0.4 (0.9) |
| WTS37P | 2.2 ± 1.0 (1.2) |
| MAXS37P | 2.2 ± 1.4 (1.2) |
| WTG48L | 14.4± 0.5 (8.1) |
| MAXG48L | 14.4± 1.4 (8.1) |
| WTQ58E | 4.6± 2.5 (2.6) |
| MAXQ58E | 5.6± 3.8 (3.2) |
| WTI84V | 1.3 ± 0.9 (0.7) |
| MAXI84V | 1.5± 0.8 (0.8) |
| WTL89I | 1.5± 0.5 (0.8) |
| MAXL89I | 2.1± 0.6 (1.2) |

<sup>1</sup>Fold change.

To explore the favored emergence in the presence of ATV of G48L, Q58E, I84V and I89L in the MAX background and to determine the effect of these substitutions in the WT background, we used site-directed mutagenesis to introduce this these mutations in both the WT and MAX protease backgrounds. The substitution S37P, preferentially selected in the MAX background when the virus was propagated in the absence of drug (Table 2), was also included in this analysis. We found that the five generated mutants displayed comparable IC_50_ values to ATV in both backgrounds, WT and MAX (Table 3). We also determined the replication capacity of these two ten mutant viruses in MT-4 cells with and without ATV (Fig. 2). Similar to the IC50 results, the five tested variants exhibited similar replication capacities in both backgrounds, WT and MAX, in either absence or presence of 20nM ATV (Fig. 2). Only WTG48L and MAXG48L showed a lower replication capacity in the absence of drug. Likewise, only substitutions at positions G48L and Q58E conferred an advantage when viruses were propagated in the presence of 20nM ATV. These results demonstrated that the G48L, Q58E, I84V and I89L substitutions were not intrinsically prohibited in the WT protease background, and that other factors must explain their low *in vivo* frequency.

The amino acid mutant repertoire also differed between the WT and MAX viruses when they were propagated without drugs (Table 2). Only one variant, D30N, was detected in both virus populations. We do not know whether the observed variants are adaptive or neutral mutations. Regardless, completely different mutant spectra were detected in these two viruses.

We performed a maximum likelihood phylogenetic reconstruction of all WT and MAX unique amino acid variants that were recovered after 32 MT-4 cell passages in the presence of ATV or DRV. Remarkably, the results showed that the two viruses, which shared an identical starting amino acid sequence, followed different evolutionary trajectories (Fig. 3). Upon visual inspection of these phylogenetic trees, it was also apparent that the MAX protease generated higher amino acid variant diversity (see below).

**FIG 3.**
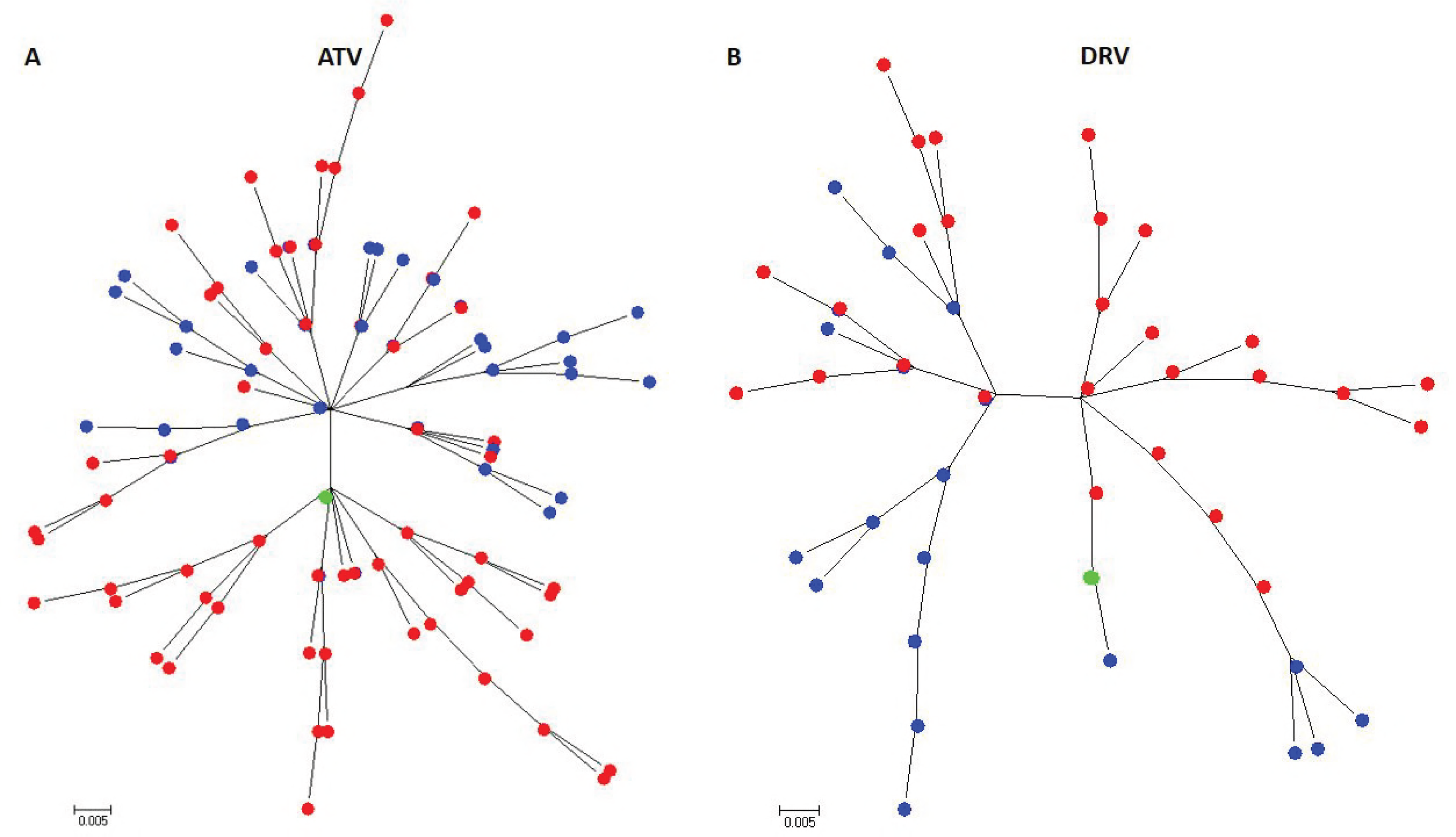
Maximum likelihood phylogram of wild-type (WT) and MAX unique HIV-1 protease amino acid variants selected after 32 passages in MT-4 cells and in the presence of **(A)** atazanavir (ATV) or **(B)** darunavir (DRV). Phylogenetic reconstruction was generated using a Jones-Taylor Thornton (JTT) model as implemented in the MEGA6 software package. Both phylogenetic trees showed that the WT and MAX viruses, which shared an identical starting amino acid sequence, followed different evolutionary trajectories. Blue and red labels correspond to WT and MAX variants, respectively. Green labels represent the starting HXB2 protease amino acid sequence.

We next compared the overall population nucleotide diversity of the WT and MAX proteases after 32 passages in the absence or presence of ATV or DRV (Table 4). Overall, nucleotide sequence diversity was significantly higher in MAX populations propagated with ATV, 0.00946 ± 0.00005 vs. 0.00517 ± 0.00004 (*P* < 0.0001), but not meaningful differences were observed in the presence of DRV, 0.00490 ± 0.00004 vs. 0.00482 ± 0.00003. As expected, in the presence of a PI, we detected a higher number of nonsynonymous substitutions than synonymous substitutions in both the WT and MAX virus proteases. However, the MAX populations always displayed a significantly higher diversity (*P* < 0.0001) of either synonymous or nonsynonymous mutations (Table 4). Diversity was also strikingly higher in MAX populations when viruses were propagated in the absence of drug (0.00095 ± 0.00001 vs. 0.00057 ± 0.00001, *P* < 0.0001) (Table 4). Interestingly, in the absence of drugs, the MAX population showed significantly higher nonsynonymous diversity (0.00117 ± 0.00001 vs. 0.00061 ± 0.00001, *P* < 0.0001) but not synonymous diversity (0.000310 ± 0.00002 vs. 0.00053 ± 0.00002, *P* < 0.0001). This suggested that the WT and MAX viruses are subjected to different selective forces in the absence of pressure from a PI. Compared to WT, the MAX populations also showed higher Shannon’s entropy values, another parameter for measuring genetic population diversity (Table 4). Notably, after 32 passages in cell culture, we observed no significant reversions of the starting synonymous substitutions introduced in the MAX protease, either in the presence or absence of PIs. Overall, our results demonstrated that MAX viruses displayed higher population genetic diversity after 32 passages in cell culture in the presence or absence of PIs.

**TABLE 4.** Summary of population metrics

| Protease | Bases<br>sequenced | Mutations<br>detected (%) | Sequence<br>diversity<br>(p-distance) | Synonymous<br>diversity<br>(p-distance) | Nonsynonymous<br>diversity<br>(p-distance) | Sn<sup>1</sup> |
| --- | --- | --- | --- | --- | --- | --- |
| WTP1 | $8.8 \times 10^6$ | 2691 (0.03) | $0.00047 \pm 0.00000$ | 0 | $0.00062 \pm 0.00001$ | 0.04 |
| MAXp1 | $8.5 \times 10^6$ | 4257 (0.05) | $0.00085 \pm 0.00004$ | 0 | $0.00113 \pm 0.00006$ | 0.05 |
| WTP32 | $7.2 \times 10^6$ | 2465 (0.03) | $0.00057 \pm 0.00001$ | $0.00053 \pm 0.00002$ | $0.00061 \pm 0.00001$ | 0.05 |
| MAXp32 | $5.2 \times 10^6$ | 2924 (0.06) | $0.00095 \pm 0.00001$ | $0.000310 \pm 0.00002$ | $0.00117 \pm 0.00001$ | 0.06 |
| WTP32 ATV | $1.9 \times 10^7$ | 71206 (0.37) | $0.00517 \pm 0.00004$ | $0.00016 \pm 0.00001$ | $0.0067 \pm 0.00003$ | 0.2 |
| MAXp32 ATV | $2.5 \times 10^7$ | 180514 (0.73) | $0.00946 \pm 0.00005$ | $0.000356 \pm 0.00001$ | $0.01252 \pm 0.00005$ | 0.27 |
| WTP32 DRV | $4.1 \times 10^7$ | 148066 (0.36) | $0.00482 \pm 0.00003$ | $0.00016 \pm 0.00001$ | $0.00624 \pm 0.00003$ | 0.16 |
| MAXp32 DRV | $1.9 \times 10^7$ | 95202 (0.73) | $0.00490 \pm 0.00004$ | $0.00026 \pm 0.00001$ | $0.00756 \pm 0.00004$ | 0.17 |

<sup>1</sup>Shannon's entropy.

## DISCUSSION

Previous research shows that codon usage can determine the mutational robustness, evolutionary capacity, and virulence of poliovirus (32). Earlier results indicate that polioviruses with synonymously mutated capsids were less mutationally robust and displayed an attenuated phenotype in an animal model. However, that study did not focus on how synonymous mutations might affect the development of escape mutations to overcome specific selection pressure targeting a precise virus gene. Here we tested the extent to which a synonymously recoded HIV-1 protease reacted to the specific selective pressure of a PI. Our present study also explored the evolvability of a retrovirus which, in contrast to other RNA viruses, integrates into the host cell genome such that viral proteins are translated from mRNAs using host cellular machinery.

We found that the WT and MAX protease viruses displayed different patterns of resistance mutations after PI treatment. These findings extend those of Lauring et al. (32), confirming that synonymously recoded and WT HIV-1 proteases occupy different sequence spaces. We further demonstrated that although the MAX and WT proteases occupied different sequence spaces, they still showed similar development of phenotypic resistance to PIs. These findings indicate that the recoded protease did not attenuate the virus’ capability to develop PI resistance, strongly suggesting that the MAX protease was as robust as the WT protease with regards to this trait. To our knowledge, this is the first study to investigate the evolvability of a synonymously recoded virus enzyme. Even if the resistance selection was conducted in MT4 cells and the result probably would be qualitatively different in primary cells, our results build on and augment the convincing evidence that recoded proteins occupy a different sequence space.

In some instances, the different mutant repertoire within the MAX protease background can be easily explained by proximity within the corresponding sequence space (e.g., G16E and L89I). However, with regards to other mutations, the explanation for the difference is not readily apparent (e.g., S37P, G48L, Q58E and I84V). In particular, the G48L substitution is very rarely selected *in vivo* in patients undergoing PI therapy (37). However, when the above substitutions (S37P, G48L, Q58E, I84V or L89I) were introduced in the WT sequence they showed parallel replication capacities to those observed with a MAX background. We can speculate that the introduced synonymous substitutions affected neighboring residues (e.g., RNA structure). However, it must be noted that the MAX and WT proteases have similar RNA folding free energy (10).

One plausible explanation is based on epistatic interactions between protease amino acid substitutions. Epistasis is a phenomenon by which a mutation’s impact on protein stability or fitness depends on the genetic background in which it is acquired (39). Complex mutational patterns often arise during the development of resistance to HIV-1 protease inhibitors. More therapy-associated mutations accumulate under PI therapy than under all other types of antiretroviral therapy. Moreover, among patients experiencing therapy failure, the majority of *in vivo* drug-experienced protease sequences include over four mutations associated with PI therapy (40). Recent findings suggest that the consequences of acquiring primary HIV-1 protease resistance mutations depend on epistatic interactions with the sequence background (41). In our study, over 80% of the MAX protease clones harboring the G48L mutation also had the I89L substitution. As mentioned above, I89L requires two nucleotide substitutions in the WT background and only one in the MAX background. In either case, our results strongly suggest that the MAX protease’s sequence position affects its genotypic PI resistance profile. Synonymous codons differ in their propensity to mutate and, as previously suggested (25, 32, 42), this differential access to protein sequence space may affect adaptive pathways.

Another intriguing finding of our study is that the MAX virus showed higher population diversity in the recoded and targeted gene following propagation in both the absence and presence of PIs. Again, we can speculate that although the MAX virus shows high fitness in tissue culture, it is subjected to greater pressure to changing or reverting to a WT synonymous background. However, remarkably, we detected almost no reversions of the synonymous substitutions introduced in the MAX protease following propagation in the presence or absence of PIs. One limitation of our study is that we investigated only one virus enzyme or protein. Further studies should include other virus proteins and other selective pressures (e.g., neutralizing antibodies and cellular virus restriction factors).

It has been suggested that RNA virus synonymous recoding can be used to push a virus to a sequence space region having a low density of neutral mutations (32). Such a lack of access to neutral substitutions could potentially reduce the virus’ capacity to generate fit progeny and adaptability to the host’s selective pressures, such that this method might serve as a new strategy for development of attenuated vaccines. Our present data suggest that this approach must be developed cautiously, and support a need to evaluate the long-term stability of synonymously recoded viruses and to carefully test individual candidates.

## MATERIALS AND METHODS

### Cell line and viruses

MT-4 cells were obtained from the National Institutes of Health (NIH) AIDS Research and Reference Reagent Program, and were grown in Roswell Park Memorial Institute (RPMI) 1640 L-glutamine medium supplemented with 10% heat-inactivated fetal bovine serum (FBS) (Gibco). The utilized WT virus corresponded to the HIV-1 HXB2 strain (http://www.hiv.lanl.gov) (Genbank accession number: K03455). The synthetic MAX HIV-1 protease was generated by PCR amplification with a combination of three overlapping synthetic DNA oligonucleotides, as previously described (10). In MT-4 cells, the MAX protease PCR product was recombined with a protease-deleted HXB2 infectious clone that had been previously linearized with BstE II (43). The protease PCR oligonucleotides used to reconstruct the full-length protease have been previously described (43). WT and MAX S37P, G48L, Q58E, I84V and I89L mutants were generated by site-directed mutagenesis using overlap extension PCR with mutated oligonucleotides as previously described (44). Again, the mutant protease PCR products were recombined with the protease-deleted HXB2 infectious clone in MT-4 cells. Cell culture supernatants were harvested at 3, 5, and 7 days post-transfection when the HIV-1 p24 antigen concentration surpassed 500 ng/ml as measured by the Genscreen HIV-1 Ag assay (Bio-Rad). Virus titration was performed in MT-4 cells, and values were expressed as tissue culture dose for 50% infectivity (TCID_50_) as previously described (45).

### Replication capacity assays

Viral replication kinetics were analyzed by infecting 1 × 10^6^ MT-4 cells with 200 TCID_50_ (MOI of 0.0002). The infected cells were incubated for 4 h at 37°C and 5% CO_2_, washed twice with phosphate-buffered saline (PBS), and then resuspended in RPMI medium supplemented with 10% FBS. To quantify viral replication, we measured the HIV-1 capsid p24 antigen concentration in 200-μl aliquots of supernatant collected every 24 h for 4–6 days. Growth kinetics were analyzed by fitting a linear model to the log-transformed p24 data during the exponential growth phase using maximum likelihood methods as previously described (10).

### HIV-1 drug susceptibility tests

ATV and DRV were obtained from the NIH AIDS Research and Reference Reagent Program. Following virus propagation and titration, we used a tetrazolium-based colorimetric method to determine the HIV-1 drug susceptibility (IC_50_) to ATV and DRV in MT-4 cells using a MOI of 0.003, as previously described (10, 46).

### Selection of ATV- and DRV-resistant viruses

WT and MAX viruses were added at an MOI of 0.01 to 1 × 10^6^ MT-4 cells, and the cells were maintained as described above. After 4 days, we transferred one-tenth of the culture, including cells and supernatant, into 1 × 10^6^ fresh MT-4 cells. All virus passages were performed in duplicate. Virus production was monitored by measurements of p24 antigen. The starting concentrations were 4 nM ATV and 3 nM DRV. Through the passages, the drug concentration was increased until reaching 40 nM ATV and 25 nM DRV. In parallel, both viruses were also propagated without either drug.

At passages 1 and 32, 140-μl aliquots of culture supernatant were collected, from which we isolated WT and MAX viral genomic RNA using the QIAamp Viral RNA Kit (QIAGEN). This purified viral RNA was then reverse transcribed and PCR amplified using the SuperScript III First-Strand Synthesis System for RT-PCR (Invitrogen) and 10 pmol of the corresponding protease oligonucleotides, which are described elsewhere (43). Details of this protocol were previously reported (47, 48). These PCR products were the starting material for performing ultra-deep sequencing.

### Ultra-deep sequencing

Massive parallel sequencing was performed in the MiSeq (Illumina) platform. Libraries of 558-nt DNA fragments were ligated to the Illumina adapters using the KAPA HyperPrep Kit (Roche #07962347001), SeqCap Adapter Kit A (Roche #07141530001), and SeqCap Adapter Kit B (Roche #07141548001). The products were purified using KAPA Pure Beads (Roche #07983280001). All libraries were quantified using the Qubit^®^ dsDNA HS Assay Kit (ThermoFisher #Q32854) and a Qubit Fluorometer (ThermoFisher #Q33216), and were qualified using the Agilent DNA 1000 Kit (Agilent #5067-1504) and a bioanalyzer (Agilent #G2939BA). Sequencing was performed using the Illumina MiSeq^®^ Reagent Kit v3 (600 cycle) (Illumina #MS-102-3003) following manufacturer’s protocol. Sequencing and paired-end analysis were performed to obtain robust fastq data for bioinformatics analysis. We obtained a mean of 50,000 sequences (reads) per amplicon and per patient sample.

Fastq files by index and pool were obtained from MiSeq and submitted to FLASH (49). The 2 × 300 paired-end reads were overlapped to reconstruct the amplicons, with the minimum number of overlapping nucleotides set to 20, and the maximum number of overlapping mismatches set to 10%. The subsequent analysis was performed as previously described (50). Briefly, fastq files were demultiplexed using amplicon oligonucleotides, and oligonucleotides were trimmed at both ends. Each amplicon and strand read was pairwise aligned with respect to the reference WT sequence, insertions were removed, and deletions were repaired if fewer than three gaps were produced. Reads with multiple indeterminations were removed, while reads having a single indetermination were repaired as per the reference sequence. Filtered and repaired reads were collapsed into haplotypes with corresponding frequencies. Haplotypes with abundances below 0.1% or that were unique to the forward or reverse strands were removed. Haplotypes common to the forward and reverse strands and with abundances of ≥0.1%, were considered consensus haplotypes, and their frequencies were summed. In the final step to remove artifacts, consensus haplotypes with abundances below 0.5% were filtered out. All computations were performed in the R language and platform, using in-house developed scripts as well as the packages Biostrings (R package 2.24.1, 2012), Ape (51), and Seqinr (52).

Virus population genetic diversity (p-distance) was determined using the MEGA6 software package (53). To determine possible selective pressures, the MEGA6 software package was used to calculate the proportion of synonymous substitutions per potential synonymous sites, and the proportion of nonsynonymous substitutions per potential nonsynonymous sites. Shannon’s entropy values were calculated as Sn = −Σ_i_ (p_i_ln p_i_)/ln N, where N is the total number of analyzed sequences and p_i_ is the frequency of each sequence in the viral quasispecies. Sn values vary from 0 (no complexity) to 1 (maximum complexity) (54). The phylogenetic reconstructions were also performed using the MEGA6 software package.

### Statistical analysis

Virus population diversity was compared by unpaired t-test using GraphPad Prism version 7 for Windows. The significance of the difference between replication kinetic slopes and IC_50_s was calculated using an unpaired t-test with Whech’s correction as implemented in GraphPad Prism.

## ACKNOWLEDGEMENTS

This study was supported by the Spanish Ministry of Economy and Competitiveness (SAF2016-75277-R) by Instituto de Salud Carlos III (PI16/00337), and cofinanced by the European Regional Development Fund (ERDF). MN was supported by the Instituto de Salud Carlos III through the Spanish AIDS network (RD16/0025/0041). AJ-P was supported by a contract of the Spanish Ministry of Economy and Competitiveness (BES-2014-069931).

